# Population connective field modeling reveals proto-retinotopic visual cortex organization in the prenatal human brain

**DOI:** 10.1101/2024.07.08.602507

**Authors:** So-Hyeon Yoo, Anne-Sophie Kieslinger, Michael A. Skeide

## Abstract

The visual space is sampled by cortical field maps in which nearby neuronal populations encode nearby locations of images received from the retina. Whether retinotopic cortical organization already emerges in the neonatal or even prenatal human brain is currently unknown. To answer this question in vivo, we applied population connective field modeling to 871 resting-state functional magnetic resonance imaging datasets ranging from prenatal to young adult age. We found topographically organized eccentricity and polar angle connectivity maps in V2 and V3 of the visual cortex as early as the 21st week of gestation. These results highlight that human proto-retinotopic cortical maps develop in the second trimester of pregnancy, predating visual experience.

## Introduction

Visual cortices transform input signals from the retina into internal representations of the visual space providing the foundation for perceiving and interpreting environments ^1^ . These cortices are organized retinotopically, such that nearby locations of the retinal image are encoded by nearby neuronal populations ^2,3^ . In retinotopic maps, nearby neuronal populations encode eccentricity, that is, the distance from the center of the visual space. Specifically, the cortical surface area representing visual space decreases with increasing eccentricity towards anteromedial visual cortices ^4–7^ . Nearby neuronal populations also encode polar angle, that is, the angular distance between a location in visual space relative to an axis, like the vertical meridian. Polar angle progression in the opposite direction at the boundary between subregions of the visual cortex is considered a critical feature of retinotopic organization ^5–7^ .

While retinotopic visual cortex organization has been extensively studied in the adult brain, its emergence in the developing brain remains poorly understood ^1^ . Functional magnetic resonance imaging (fMRI) studies show that eccentricity and polar angle maps are in part qualitatively similar to adults by age 5 except that children have a similar amount of V1 surface area representing the lower and upper vertical meridians of the visual space ^8–10^ . In macaque monkeys, however, retinotopically organized eccentricity and polar angle maps have been identified already at the neonatal stage using fMR I ^11^ . Other proto-retinotopic features, including response preferences for horizontal and vertical orientation as well as high and low spatial frequencies have been revealed by fMRI in infants aged 5 months ^12^ . Whether proto-retinotopic organization is present in neonatal, let alone prenatal humans, is currently unknown.

To answer this question, we applied population connective field modeling to 871 resting-state fMRI datasets from the Human Connectome Project including prenatal, neonatal, adolescent, and adult age group s ^13, 14^ . Population connective field modeling serves to estimate the dependence between hemodynamic signals in distinct cortical regions by predicting responses in a region as a function of activity in another region based on a Gaussian function ^15^ . This method has been validated against the widely used population receptive field modeling approach using task-based fMRI signals evoked by visual stimuli ^15^ . In addition, this method has also been validated against task-free resting-state fMRI signals in V1, V2 and V3 ^16^ . Accordingly, population connective field modeling of resting-state fMRI data allowed to examine proto-retinotopic organization in prenatal subjects who are not able to engage in a visual field mapping task. Here, we used a retinotopy template that was derived from task-based fMRI data collected in adults during visual receptive field mapping experiments. Warping this template into native space we identified the regions of interest V1, V2 and V3, then employed population connective field modeling to find resting-state hemodynamic time series in V1 that correlated most strongly with corresponding time series in V2 and V3, and finally inferred eccentricity and polar angle values in V2 and V3 from the values in V1 given by the template ^15–17^ . Differences in data quality across datasets were adjusted with an empirical Bayesian framework for group comparison analyses. This framework also accounted for possible site effects on data acquisition, for example, related to hardware, software or human factors in different scanning environments.

Given that retinotopic maps have been identified in newborn macaques ^11^, we hypothesized that topographically organized eccentricity and polar angle maps in humans can be identified only from a neonatal age onwards. Contrary to our hypothesis, however, we found topographically organized eccentricity and polar angle connectivity maps already from prenatal age onwards, as early as in the 21st week of gestation.

## Results

### Group characteristics

Prenatal MRI scans recorded during the second trimester of pregnancy (here between 21 and 28 weeks gestational age) and the third trimester of pregnancy (≥29 weeks gestational age) were compared against preterm neonatal data (born <37 weeks gestational age) and full-term neonatal data (born ≥37 weeks gestational age) as well as MRI scans recorded during adolescence (12–16 years) and early adulthood (18–21 years). All six groups include cross-sectional data of different subjects (Table 1). Preterm neonatal data were not excluded since we aspired to examine whether proto-retinotopic maps would develop typically in this population which is at risk for neurodevelopmental disorders ^18,19^ .

**Table 1.** Group characteristics.

|  | **tri-2<sup>2</sup>**<br>**prenatal** | **tri-3<sup>3</sup>**<br>**prenatal** | **preterm**<br>**neonatal** | **full-term**<br>**neonatal** | **adolescent** | **adult** |
| --- | --- | --- | --- | --- | --- | --- |
| mean scan age | 26.10 | 32.00<sup>4</sup> | 38.23<sup>4</sup> | 41.29<sup>4</sup> | 14.11<sup>5</sup> | 20.21<sup>5</sup> |
| scan age range | 21–28 | 29–36<sup>4</sup> | 34–44<sup>4</sup> | 37–44<sup>4</sup> | 12–16<sup>5</sup> | 18–21<sup>5</sup> |
| sex (f/m)<sup>1</sup> | 55/54 | 41/55 | 44/68 | 203/237 | 37/23 | 32/22 |
| n subjects | 109 | 96 | 112 | 440 | 60 | 54 |
| n excluded | – | – | 2 | 19 | – | – |

<sup>1</sup>female/male <sup>2/3</sup>second/third trimester of pregnancy <sup>4</sup>age in weeks <sup>5</sup>age in years

### Controlling for anatomical differences, data quality and site effects

To account for possible anatomical effects of brain size and cortical morphology, such as gyrification and sulcal patterns, on registration quality, a step-wise transformation procedure was applied to register structural images to an adult surface template containing both the dorsal and ventral visual regions of interest V1, V2 and V3 (Supplementary Fig. S1; see Methods for details). Individual examples demonstrating image registration results across different age groups are provided in Supplementary Fig. S2.

Temporal and slice signal-to-noise ratio (tSNR and sSNR) in V1–V3 differed significantly between the six groups (tSNR: χ ^2^ (5) = 505.96, η^2^ = 0.76, bootstrapped p < 0.001, Kruskal-Wallis test; sSNR: χ ^2^ (5) = 179.73, η^2^ = 0.45, bootstrapped p < 0.001, Kruskal-Wallis test) (see Supplementary Fig. S3 for pairwise group comparisons). One possible factor among others contributing to this effect could be that prenatal and neonatal images were acquired with shorter repetition times compared to adolescent and adults. Accordingly, to account for these differences in data quality and also for data acquisition site effects in direct statistical group comparisons, eccentricity and polar angle values obtained from the population connective field model were adjusted for tSNR values and sites by using a parametric empirical Bayesian framework ^20^ . This framework was neither used to put less weight on individuals with low tSNR nor to adjust model fits but to remove the effects of tSNR from the eccentricity and polar angle values before determining statistical group differences.

### Population connective field modeling

Hemodynamic time courses estimated by the population connective field model in V1 of the visual cortex were first correlated with observed hemodynamic time courses measured in V2 and V3. Eccentricity and polar angle values were then obtained from a template that was derived from event-related fMRI data collected in adults during visual receptive field mapping experiments (Fig. 1 and 2). To this end, template values at each vertex in V1 were projected to the vertex in V2 and V3 that was fit best by the model as indicated by the highest positive correlation coefficient. The variance explained differed significantly between groups (Left: χ ^2^ (5) = 43718.80, η^2^ = 0.10, bootstrapped p < 0.001, Right: χ2(5) = 26646.76, η^2^ = 0.07, bootstrapped p < 0.001, Kruskal-Wallis test), driven by a gradual increase in model fits from prenatal to adult age (from mean R ^2^ = 0.09 to 0.12) . Finally, we confirmed statistically that associations between the connective field model and the retinotopic template data did not arise by chance. Accordingly, eccentricity and polar angle values obtained from the empirical hemodynamic signal were compared with values obtained from simulated random hemodynamic signals (Supplementary Fig. S8 and S11). Connective field sizes differed significantly between groups (Left: χ ^2^ (5) = 92098.34, η^2^ = 0.15, bootstrapped p < 0.001, Right: χ2(5) = 123467.50, η^2^ = 0.16, bootstrapped p < 0.001, Kruskal-Wallis test) and the increase in connective field size across eccentricity became steeper with age (see Supplementary Fig. S4). The variance explained based on split-half cross-validation is shown in Supplementary Fig. S5.

**Fig. 1.**
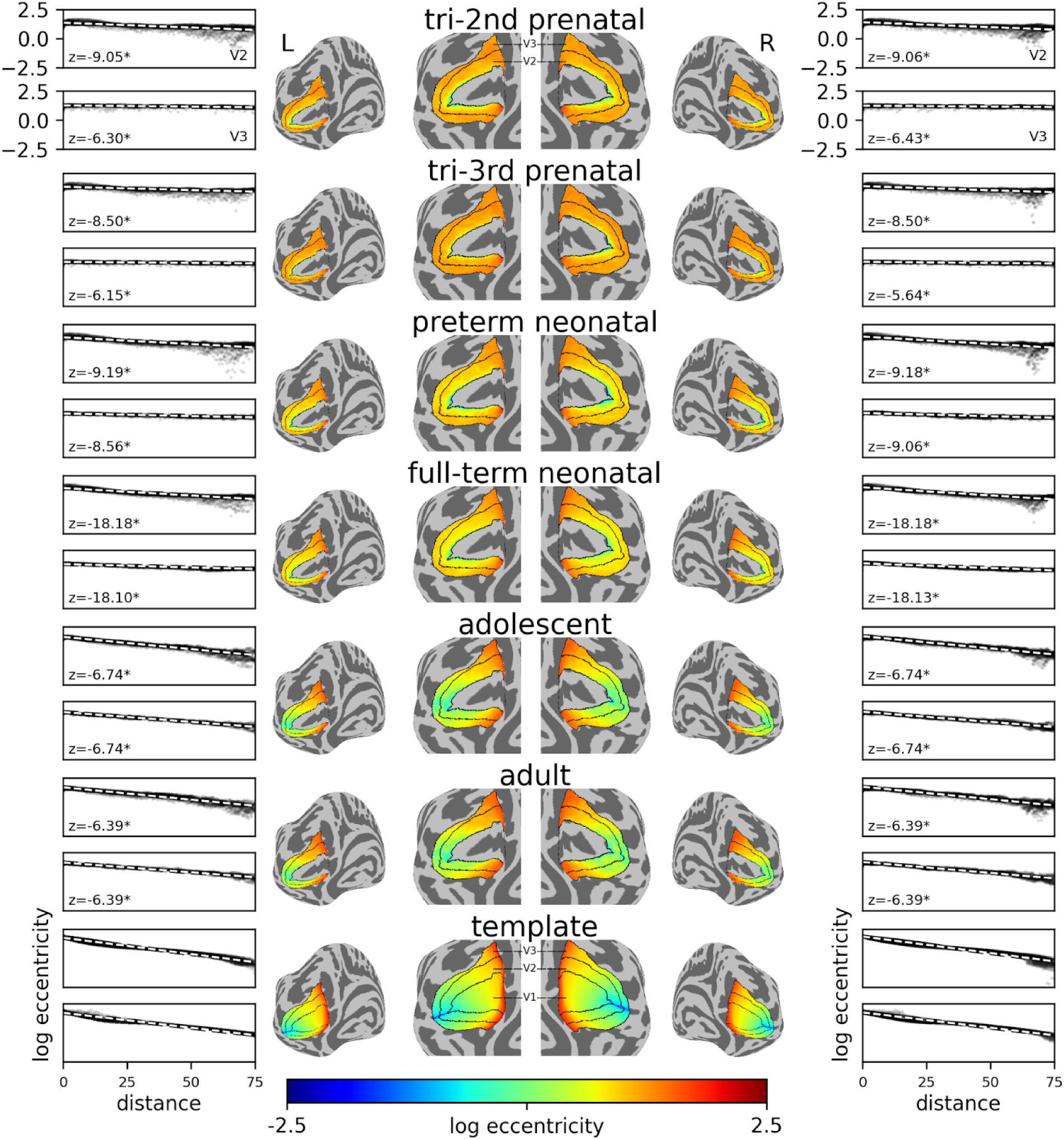
Eccentricity maps. Eccentricity values (degrees of visual angle) were obtained from a template (shown in the bottom row) generated using event-related fMRI data collected in a receptive field mapping experiment. Maps were averaged within each group. L = left hemisphere, R = right hemisphere. Black solid lines demarcate V1, V2, and V3 on the cortical surface. . Diagrams summarize log-transformed eccentricity (y-axis) and distance (mm) from the medial edges of V2 and V3 (x-axis) across all subjects and separately for each region of interest and hemisphere. Dashed lines are linear regression fits. In all groups, individual slopes of the linear regression differed significantly from zero at a threshold of p < 0.01 (Wilcoxon signed-rank tests) confirming statistically that eccentricity decreased with distance from the medial edge (marked by an asterisk). These analyses were not run on the template as it lacks individual slopes.

**Fig. 2.**
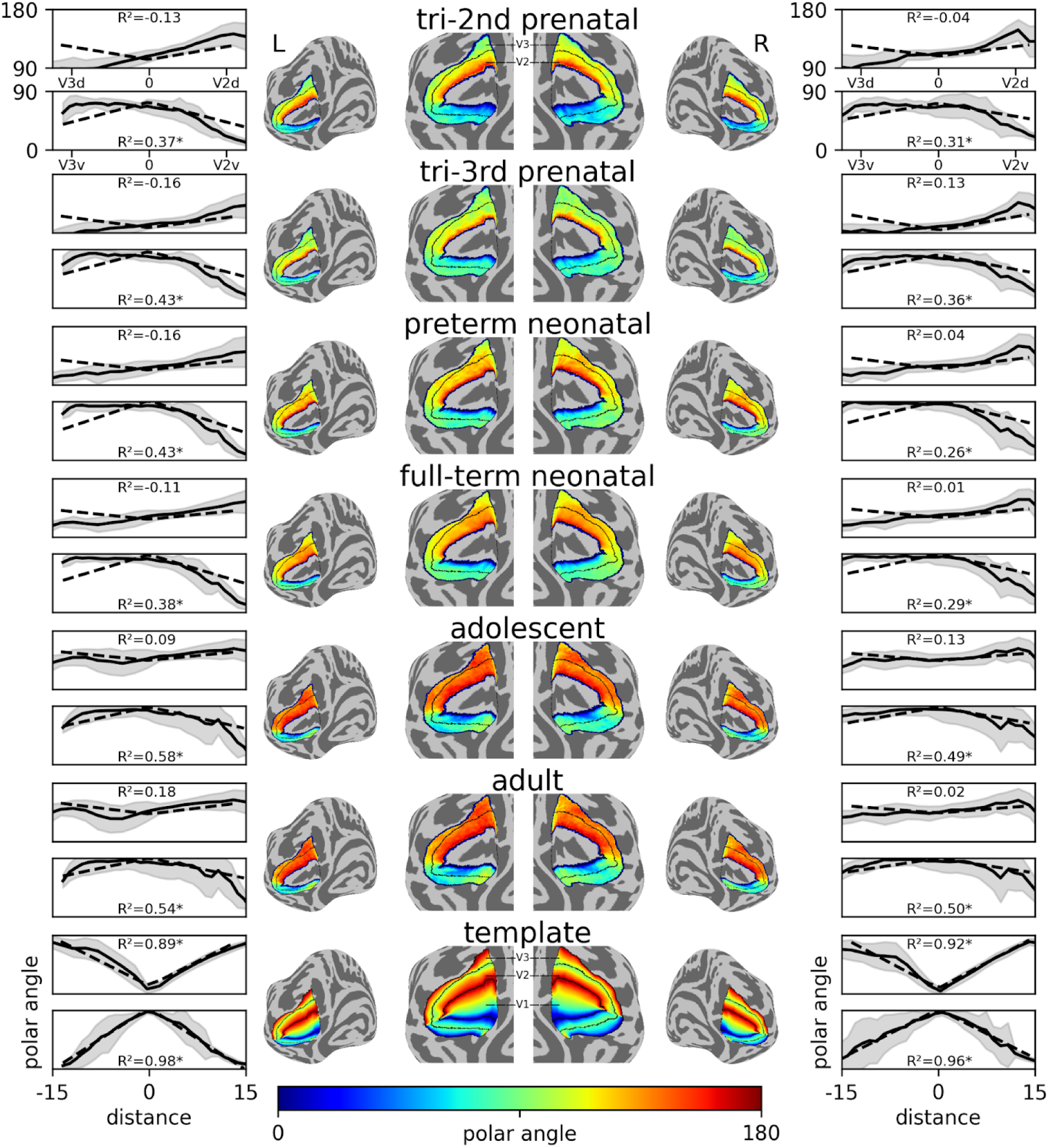
Polar angle maps. Polar angle values (degrees of visual angle) were obtained from a template (shown in the bottom row) generated using event-related fMRI data collected in a receptive field mapping experiment. Maps were averaged within each group. L = left hemisphere, R = right hemisphere. Black solid lines demarcate V1, V2, and V3 on the cortical surface. Diagrams summarize polar angle (y-axis) and distance (mm) from the V2-V3 boundary (x-axis) across all subjects and separately for each region of interest and hemisphere. Solid lines represent averaged polar angle values and shaded areas represent the distribution density of polar angle values across distance. Dashed lines are fits to an absolute value function. Asterisks indicate statistically significant fits at a threshold of p < 0.01 (absolute value regression).

### Eccentricity connectivity mapping

Mapping log-transformed eccentricity values onto the surfaces revealed steady decreases with distance from the medial edges of V2 and V3 in all groups (Fig. 1). Moreover, we confirmed statistically that progression slopes differed significantly from zero, the equivalent of no change of eccentricity over distance, in all groups (all z ≤ 5.36, all p < 0.01, Wilcoxon signed-rank tests). At the same time, we found significant group differences between eccentricity maps and the number of significant vertices gradually increased with increasing age differences (see Supplementary Fig. S6). Examples of individual eccentricity maps across the different age groups are provided in Supplementary Fig. S7.

Eccentricity maps without predefined extrastriate regions of interest can be found in Supplementary Fig. S8. In all groups and in both hemispheres, the connective field model applied to the empirical data fit the eccentricity template better than simulated data, as indicated by comparing mean squared differences (all z ≥ 6.39, all p < 0.001, Wilcoxon signed-rank tests; see Supplementary Fig. S9 for distributions of mean squared differences across all groups).

### Polar angle connectivity mapping

Modeling the v-shaped distribution of polar angle values by fitting absolute value functions we found that, in all groups, the progression of polar angle reversed direction at the ventral but not the dorsal boundary between V2 and V3(Fig. 2). Specifically, in the ventral part of V2 and V3, the model explained a considerable amount of variance increasing from R² = 0.37 in the second trimester of pregnancy to R² = 0.82 in adulthood while fits were substantially lower in the dorsal part of V2 and V3 (max. R² = 0.14). This discrepancy is in line with prior work consistently reporting higher spatial variability in dorsal compared to ventral visual cortex ^17,21^ . To quantitatively confirm polar angle reversal we also computed gradients representing the direction of change within vertex-wise polar angle values. Mean gradients in a dorsal-to-ventral coordinate system indicated that polar angle in V2 and V3 indeed progressed in opposite directions in all groups (Fig. 3). There were also significant group differences between polar angle maps and the number of significant vertices gradually increased with increasing age differences (see Supplementary Fig. S10). Examples of individual polar angle maps across the different age groups can be found in Supplementary Fig. S11. Polar angle maps without predefined extrastriate regions of interest can be found in Supplementary Fig. S8. In all groups and both hemispheres, the connective field model applied to the empirical data fit the polar angle template better than the simulated data (all z ≥ 6.39, all p < 0.001, Wilcoxon signed-rank tests; see Supplementary Fig. S12 for distributions of mean squared differences across all groups).

**Fig. 3.**
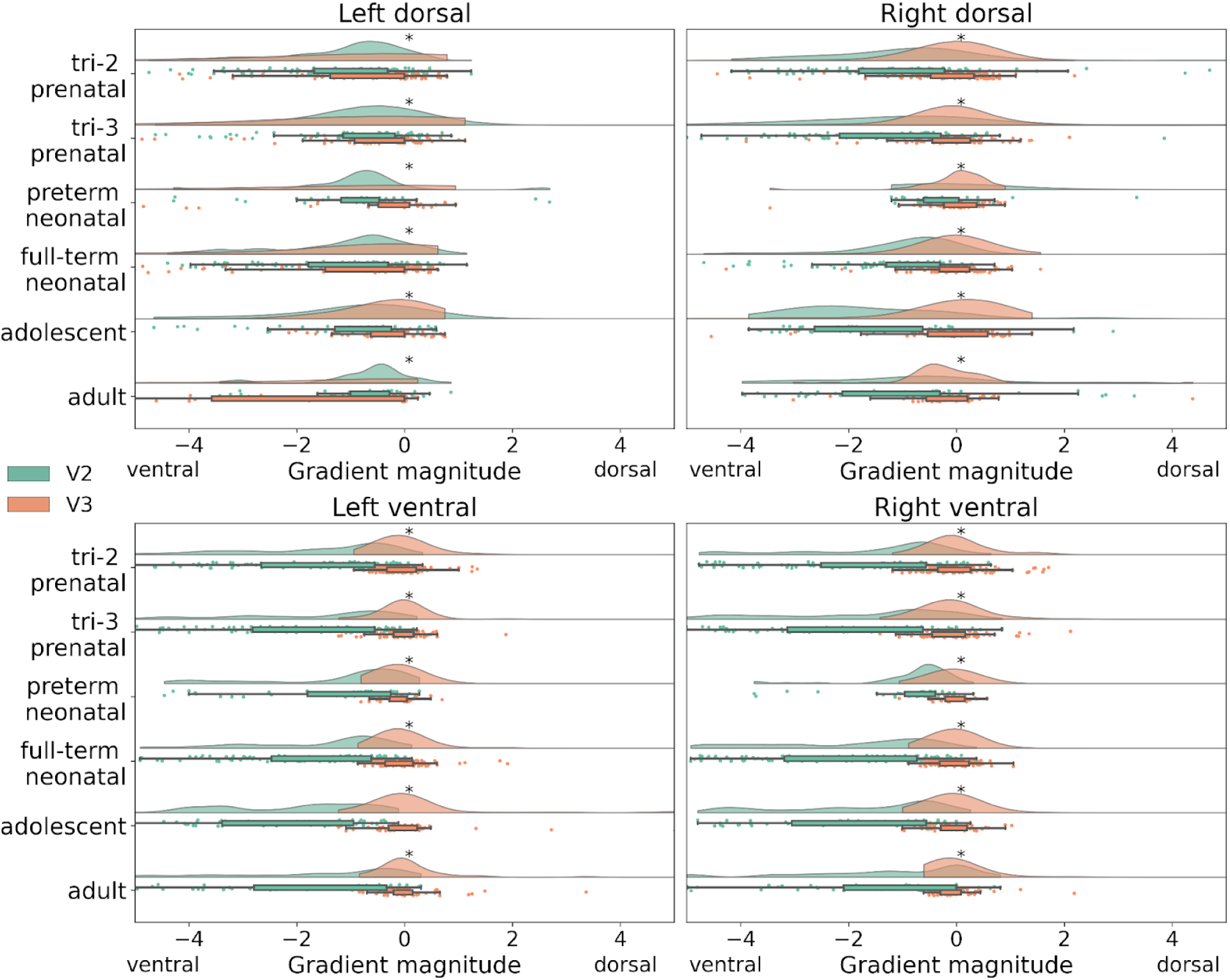
Mean gradient directions of polar angle values. Mean gradient magnitudes were computed to quantify dorsal and ventral directions of polar angle progression. To this end, V2 (green) and V3 (orange) were partitioned into dorsal and ventral subregions. x-axes depict the magnitude of the gradients indicating the direction of increasing values and y-axes depict the different age groups. Negative values indicate ventral increases while positive values indicate dorsal increases . Curves illustrate probability density while lines illustrate the lengths of the distributions. Box plots visualize upper and lower quartiles. Asterisks indicate statistically significant differences at a bootstrapped threshold of p < 0.001 (Wilcoxon signed-rank tests).

## Discussion

Population connective field modeling of developmental resting-state fMRI data revealed a proto-retinotopic organization of visual cortices in prenatal human brains. Already in the 21st gestational week, eccentricity decreased with distance from the medial edge of V2 and V3and polar angle reversed directions at the ventral border of V2 and V3.

Human visual population receptive fields are traditionally mapped using task-based fMRI. This technique, however, is not applicable to prenatal subjects and has not yet been successfully applied to neonatal subjects. Accordingly, population connective field modeling was employed in this study to reconstruct retinotopic features from resting-state fMRI data in the absence of a visual task. The task-free resting-state fMRI results reported here can be explained by animal models based on the observation that temporally coupled stimulus-independent neural activity is necessary for the development of visual receptive field properties related to retinotopic map s ^22–25^ . Such spontaneous activity is thought to strengthen or maintain correctly targeted synaptic connections. Synchronized stimulus-independent firing can be observed in primate and rodent visual cortex well before behavioral signs of visual processing emerge and even before photoreceptors respond to light ^22–25^ . Future studies using longitudinal high-field MRI might be able to shed light on these processes in vivo and noninvasively at a mesoscopic scale capturing layer-specific hemodynamic responses and fiber projections.

In addition to spontaneous cortical activity, retinal waves of ganglion cell activity during gestation are thought to serve to establish retinotopic visual cortex organization in the prenatal vertebrate nervous system. In line with this prediction, retinal waves have been observed well before birth in a number of species including nonhuman primates, cats, mustelidae and rodents ^26–28^ . It can be expected that gestational ganglion cell activity emerges at an early prenatal age in humans but this has yet to be demonstrated empirically.

Proto-retinotopic organization of eccentricity and polar angle maps in the second trimester of pregnancy is remarkable from a behavioral perspective since the eyes typically open only by 28 weeks of gestation, pupils constrict and dilate only at around 31 weeks of gestation, and the fetus orients to external laser light stimuli only by around 34 weeks of gestation ^29,30^ . Furthermore, from an environmental perspective, biophysical modeling results suggest that intrauterine illumination is severely limited by maternal tissue ^31^ . There is fMRI evidence, however, that resting-state retinotopic organization emerges without visual experience even in congenitally blind adults ^16^ . Furthermore, these results in humans are supported by single-cell recording and photon imaging studies in rodents. These studies show that, already at eye opening, the proportion of receptive fields and the angle of visual space they represent is similar to more mature animals ^32^ . At the same time, despite these similarities, neural responses became more reliable in the course of further brain development ^32^ . Such maturation effects could be related to the present finding that age group differences between eccentricity and polar angle maps gradually increased with increasing age differences.

The connective field mapping approach taken here requires a flattened cortical surface template that allows to predict the location and topographic organization of visual areas from cortical anatomy alone. Such templates are currently only available for eccentricity and polar angle ^17^ . Future work should consider applying population connective field modeling to additional retinotopic features including orientation tuning and spatial frequency tuning if they become available^12^ . Another aspect to be addressed is that the retinotopic feature templates used for the present study are derived from task-based fMRI in adults. It is an open question whether such templates could also be derived from neonatal infants despite the considerable challenges inherent to task-based fMRI in this population.

To account for differences in data quality and site effects data were adjusted with an empirical Bayesian framework for direct statistical comparison across prenatal, neonatal, adolescent and adult groups. In the other analyses, we desisted from adjustment to avoid introducing biases into the population connective field model estimation. This decision is based on previous work demonstrating that connective field size estimates are unbiased statistics which are robust to differences in signal-to-noise ratio ^16^ . Contributions of other factors beyond our analytical control, such as more variable signals in the younger developmental groups, remain to be ruled out in future investigations ^33,34^ .

## Methods

### Participants

#### Prenatal and neonatal participants

Prenatal and neonatal participants were recruited between 2015 and 2019 at St Thomas’ Hospital in London for the Developing Human Connectome Project. Written informed consent of all legal guardians was obtained for all participants prior to taking part in the study. All study procedures were reviewed and approved by the London-Riverside Research Ethics Committee of the Health Research Agency (REC: 14/Lo/1169) ^14,35^. The analysis of the pre-existing data was approved by the Ethics Committee at the Medical Faculty of the University of Leipzig, Germany (IRB00001750).

109 unique prenatal datasets corresponding to the second trimester of pregnancy (between 21 and 28 weeks gestational age) and 96 unique prenatal datasets corresponding to the third trimester of pregnancy (≥29 weeks gestational age) were included in the current study (mean gestational age 32.00 weeks, range 29–36 weeks). Additionally, we included 41 preterm neonates born <37 weeks gestational age (mean gestational age 32.46 weeks, range 23–36 weeks). 27 neonates admitted to neonatal intensive care, nine neonates with incomplete or abandoned scans, one neonate with vermian hypoplasia and one neonate with congenital heart disease were excluded from the study. The final cohort of 111 full-term neonates born ≥37 weeks gestational age (mean gestational age 39.98 weeks, range 37–42 weeks) had normal findings on clinical examination by a neonatologist prior to MRI data acquisition. Neuroradiological inspection of the MRI data mostly revealed clinically insignificant incidental finding s ^36^ . Regardless of a potential clinical significance, incidental findings with unlikely measurable relevance for functional connectivity analysis were permissible ^37^ . However, in line with the guidelines of the Developing Human Connectome Project, we excluded datasets with incidental findings that might have a possible impact on the analyses in accordance with radiological ratings.

#### Adolescent and adult participants

Adolescent and adult participants were recruited between 2017 and 2019 in Boston, Los Angeles, Minneapolis and St. Louis for the Human Connectome Project – Development. Written informed consent of all legal guardians was obtained for all participants prior to taking part in the study. All study procedures were reviewed and approved by a central institutional review board at Washington University in St. Louis. The analysis of the pre-existing data was approved by the Ethics Committee at the Medical Faculty of the University of Leipzig, Germany (IRB00001750).

60 unique adolescent datasets (mean scan age 14.49 years, range 12–17 years) and 54 unique early adult datasets (mean scan age 20.21 years, range 18–21 years) were included in the current study. 289 datasets that did not pass the quality control procedure were excluded from our analysis (see Data Preprocessing below). We also excluded 34 subjects with visual impairments. Neuroradiological inspection of the MRI data revealed no clinically significant incidental findings

### Data acquisition

#### Prenatal and neonatal data

Prenatal and neonatal images were collected on a 3T Philips Achieva scanner at the Evelina London Children’s Hospital. These data are publicly available as part of the fourth release of the Developing Human Connectome Project (https://biomedia.github.io/dHCP-release-notes/) and were accessed in the National Institute of Mental Health Data Archive (NDA) ^14^ . Example images can be found in the data release paper published by the Developing Human Connectome research group ^14^ .

Data were acquired without sedation. Neonates were scanned during natural sleep following feeding and swaddling in a vacuum-evacuated blanket. To standardize pose, subjects were placed in a cradle. Hearing protection was provided in the form of molded dental putty, Minimuffs on-ear noise attenuators, and an acoustic hood ^37^ .

For structural imaging, a T2-weighted fast spin echo image was acquired twice with TR = 12 s, TE = 156 ms, SENSE factor: axial = 2.11 and sagittal = 2.58, 290 × 290 × 203 voxels, voxel size 0.8 mm x 0.8 mm x 1.6 mm (overlapped by 0.8 mm). In addition, a T1-weighted inversion recovery fast spin echo image was acquired twice with TR = 4,795 ms, TE = 8.7 ms, SENSE factor: axial = 2.26 and sagittal = 2.66 and matched to the T2-weighted image in terms of number of voxels and voxel size ^38^ .

For prenatal functional imaging, 350 T2*-weighted single-shot echo-planar imaging volumes were acquired with TR = 2,200 ms, TE = 60 ms, multiband acceleration factor of 3, slice matrix 144 × 144, voxel size 2.2 mm x 2.2 mm x 2.2 mm ^39^ .For neonatal functional imaging, 2,300 T2*-weighted single-shot echo-planar imaging volumes were acquired with TR = 392 ms, TE = 38 ms, flip angle = 34°, multiband acceleration factor of 9, 67 × 67 × 45 voxels, voxel size 2.15 mm x 2.15 mm x 2.15 mm. Single-band reference scans were also acquired with bandwidth-matched readout, along with additional spin-echo echo-planar imaging acquisitions (fieldmaps) with 4 x AP and 4 x PA phase-encoding directions ^38^ .

#### Adolescent and adult data

Adolescent and adult data were collected on 3T Siemens Prisma scanners with 32-channel head coils at Harvard University, University of California at Los Angeles, University of Minnesota, and Washington University in St. Louis. These data are publicly available as part of the second release of the Human Connectome Project – Development and were accessed in the National Institute of Mental Health Data Archive (NDA) ^13^ .

For structural imaging, a T2-weighted variable-flip-angle turbo-spin-echo image was acquired with TR = 3,200 ms, TE = 564 ms, 320 × 300 × 208 voxels, voxel size 0.8 mm x 0.8 mm x 0.8 mm). In addition, a T1-weighted multi-echo image was acquired twice with TR = 2,500 ms, TE1 = 1.81 ms, TE2 = 3.6 ms, TE3 = 5.39 ms, TE 4 = 7.18 ms and matched to the T2-weighted image in terms of number of voxels and voxel size. Subjects watched a movie during these procedures ^40^ .

For resting-state functional imaging, 488 T2*-weighted single-shot echo-planar imaging volumes were acquired four to six times with TR = 800 ms, TE = 37 ms, flip angle = 52°, multiband acceleration factor of 8, 67 × 67 × 45 voxels, voxel size 2 mm x 2 mm x 2 mm. Subjects were instructed to look at a fixation cross during this procedure. A single-band spin-echo echo-planar imaging fieldmap with one AP and one PA phase-encoding direction was also acquired with bandwidth-matched readout ^40^ .

## Data preprocessing

### Developing Human Connectome Project: Prenatal and neonatal images

#### Structural image preprocessing

T2-weighted structural images were pre-processed according to the structural pipeline developed in the developing human connectome project (dHCP) ^41^ . First, motion correction and super-resolution reconstruction techniques were employed on T1- and T2-weighted structural images ^42,43^ . Second, the reconstructed T1- and T2-weighted images were bias-corrected using the N4 bias field correction algorithm ^44^ and the Brain Extraction Tool (BET) implemented in the FSL software package (5.0.11) ^45^ . Finally, each T1-weighted image was registered rigidly to the corresponding T2-weighted image using the Medical Image Registration ToolKit (MIRTK).

After preprocessing, the T2-weighted images were segmented into different tissue types (white matter, cortical gray matter, subcortical gray matter, and cerebrospinal fluid) using the Developing brain region annotation with Expectation-Maximization (Draw-EM) algorithm ^46^ . Draw-EM was also employed to generate surface meshes (white matter, pial, and mid-thickness surfaces). Only those datasets that exhibited an identical number of vertices in the white matter and pial meshes following the reconstruction of surface data were subjected to further analysis. This step is necessary as the vertices in white matter and pial mesh form a triangular prism when resampling a volume image to a surface image. During this process, the signals of multiple voxels in the triangular prism are restructured into a single time series. Accordingly, if the number of vertices in white matter and pial meshes are different, volume-to-surface mapping fails. 95 out of 136 preterm datasets and 346 out of 457 full-term datasets with this quality issue were excluded from further analysis.

#### Functional image preprocessing

T2*-weighted functional images were preprocessed in accordance with the resting-state functional processing pipeline in the dHCP ^37^ . First, local signal distortions resulting from magnetic field inhomogeneities were corrected using the FSL TOPUP too l ^47^ . Second, slice-to-volume and rigid-body realignment were employed using the FSL EDDY tool ^48^ to correct for in- and out-of-volume head motion and dynamic distortions. Third, a high-pass filter with a 150s cutoff was applied to remove slow drifts. Finally, residual head motion, motion artifacts arising from multiband acceleration, and physiological noise were removed using a linear regression model implemented in FMRIB’s ICA-based Xnoiseifier tool (FIX, v1.066) ^49^ .

### Human Connectome Project: Adolescent and adult images

#### Structural image preprocessing

T1- and T2-weighted structural images were preprocessed according to the structural pipeline of the Human Connectome Project (HCP-D). This pipeline comprises three preprocessing stages: PreFreeSurfer, FreeSurfer and PostFreeSurfer. In PreFreeSurfer, structural images were distortion-corrected, aligned, and registered to the Montreal Neurological Institute (MNI) standard space using the FMRIB software library implemented in FSL. In FreeSurfer, structural cortical maps are reconstructed by employing the recon-all pipeline. In PostFreeSurfer, a brain mask, a cortical ribbon volume image, and cortical myelin maps are generated. Finally, the surface mesh in native space is resampled into fs-LR space ^50^ .

#### Functional image preprocessing

T2*-weighted functional images were preprocessed in accordance with the resting-state functional processing pipeline of the HCP-D ^40,50^ . First, volumes were preprocessed using FSL FLIRT to correct for head motion and dynamic distortions ^51^ . FSL FLIRT was also employed for non-linear alignment to the MNI standard space. Second, distortions in the phase encoding direction were estimated using FSL TOPUP ^47^ . Third, volumes were resampled from standard space to a surface representation consisting of 91,282 gray matter coordinates. Fourth, gray matter ribbons were determined in native space with the fslmaths tool in FSL and mapped to the gray matter coordinates surface using the Connectome Workbench tool (v1.4.2). Fifth, head motion indices and independent components were regressed out using FMRIB’s ICA-FIX (v.1.0.6.14). In the final step, images were re-registered to the cortical surfaces using MSMAll (v.1.0) ^52^ .

### Population connective field modeling analysis

#### Registration of functional images from volume to surface space

Structural surfaces in native T2-weighted image space were warped into the native volume space of the functional images using the affine transformation matrices generated by dHCP (boundary-based registration, FSL FLIRT). Subsequently, functional volumetric data were projected to the resulting surface using the ribbon-constrained mapping approach implemented in the Connectome Workbench tool (v1.5.0). Regions of interest were primary, secondary, and third visual cortices (V1, V2 and V3) defined by Benson’s anatomical template ^17^ .

#### Population connective field model estimation

Population connective field modeling has been described comprehensively by Haak and colleagues (2013) ^15^ . First, hemodynamic time series for each vertex in V1, V2, and V3 were converted to time series of percent signal change.

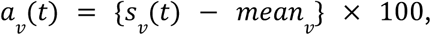

where, 𝑎 _𝑣_ (𝑡) and 𝑠 _𝑣_ (𝑡) represent the percent signal change and the raw signal of the vertex 𝑣 at time point *t*, respectively, and 𝑚𝑒𝑎𝑛_𝑣_ is a time-averaged raw signal of the vertex 𝑣.

Next, candidate connective fields were specified. The cortical connective field for each vertex 𝑣 in the target region 𝑉 2 and 𝑉 3 consists of weights for each vertex *w* in the source region 𝑉 1, with *w*_0_ denoting the center vertex of the connective field in the source region *V*1. Each candidate connective field model is a Gaussian distribution defined by its center vertex *W*_0_ and its spread over the cortical surface.

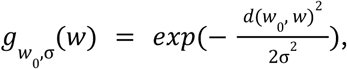

where, *d* (*w* _0_, *w*) indicates the shortest distance from the center vertex *w*_0_ to all other vertices in the source region along the cortical surface mesh and 𝑔 *_*w*_*_0_, _σ_(*w*) indicates the connective field weights for all *w*, centered on *w* _0_ with spread σ . Distances between vertices were calculated along the pial surface using a Scipy (v1.12.0) implementation of the Dijkstra algorithm in Python (v3.11.7).

For each connective field 𝑔, the weights of the cortical field in the surface mesh were then used to compute a time series. The time series 𝑎 _*w*_ (𝑡) of all vertices *w* in 𝑉1 are weighted by their contribution to the cortical field *_*w*_*_0_, _σ_(*w*) and summed up to constitute the time series of the connective field 𝑐 _*w* 0,σ_(𝑡).

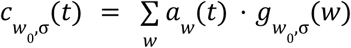

The fit of the connective field model was defined as the residual sum of squares between the time series of the connective field and the time series of the vertex in V2 and V3 𝑏 (𝑡) . For an initial parameter search, candidate time series of the connective field were calculated for all V1 vertices and 10 cortical spread values σ between 3 and 25 mm. A non-linear optimization over spread σ was performed for all vertices, using the Broyden-Fletcher-Goldfarb-Shanno algorithm implemented in Scipy’s optimization module.

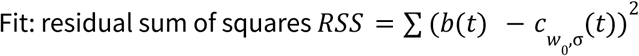

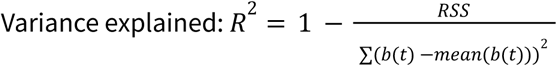

Split-half cross-validation was employed to confirm that the results obtained from connective field modeling generalize despite intrinsic noise in the data. To this end, the dataset was divided into two halves: the first half was used for model fitting, and the second half was used for testing the model. In this procedure, the model parameters, that is, the estimated spatial spread (σ) and center of the connective field, are fit using only the first half of the data, by testing a large number of candidate connective fields against the hemodynamic signal of the target vertex and then selecting the connective field with the lowest residual sum of squares. The variance explained in both datasets provides an unbiased estimate of the model’s performance and generalizability.

#### Null model estimation on simulated data

To create a null model without retinotopically organized signals, hemodynamic surface data in V1, V2 and V3 were simulated using a local correlation model. For each surface vertex, a random time series was generated from a normal distribution with the same number of timepoints as the empirically measured hemodynamic data. The local correlations in V1, V2 and V3 were then smoothed using a Gaussian filter with a kernel width of 30 mm and adjusted to match the median local correlation coefficient of the empirical data using Cholesky decomposition.

#### Retinotopic template projection

Retinotopic maps containing eccentricity and polar angle values were derived from an adult template ^17^ . As the template was defined on the fsaverage surface mesh template in FreeSurfer, it was necessary to warp the template from fsaverage space to native space. Due to the structural differences between developing and adult brains, a direct transformation from fsaverage to native space could introduce misalignment or distortions. Accordingly, a four-step registration procedure was applied to the retinotopy templates instead, including (1) registration from fsaverage space to the space of the young adult template of the Human Connectome Project (HCP), (2) registration from young adult space to the space of the 40-week template of the dHCP ^44^ (see https://t.ly/AFwNT for both hemispheres), (3) registration from 40-week space to the space of an age-matched group template, and (4) registration from the age-matched group template space to native space.

For the first registration step, we used the registration sphere of the HCP young adult mesh deformed to fsaverage, available from the HCP ^49^ . For the second registration step, we used an existing sphere generated during the transformation of the dHCP 40-week template sphere to the HCP sphere ^35^ . For the third registration, we used an existing sphere that was generated in the dHCP by transforming an age-matched template sphere to the 40-week template sphere. For the fourth registration, the native surface sphere was rotated to align with HCP fsLR volumetric space and then registered to the age-matched template sphere of the dHCP using the Biomedical Imaging Analysis Group’s Medical Image Restoration Toolkit (MIRTK) and the newMSM software package (v.0.5.1-BETA), following the procedure implemented in https://github.com/ecr05/dHCP_template_alignment. As a final step, these four registrations were concatenated and used to resample the template from fsaverage space to native space. The transformation and concatenation of all registrations was performed using the Human Connectome Project workbench toolkit (v1.5.0). As the preprocessed resting-state fMRI data provided by the HCP is resampled in fs-LR space, registration was performed only once from fs-LR to fsaverage space. All remaining analysis steps were identical between the HCP and dHCP data.

In each prenatal and neonatal subject, connective field modeling was conducted in native space using a warped template. In adolescent and adult subjects, connective field modeling was conducted in FreeSurfer’s fs-LR space. Subsequently, maps were warped into fsaverage space for further statistical analysis of positive correlation coefficients as well as connective field size, eccentricity, and polar angle values. Vertex-wise eccentricity and polar angle values were assigned to the most strongly positively correlated seed vertex.

#### Local gradient analysis

To quantitatively assess the direction of change in the V2 and V3 polar angle values, we computed a gradient at each vertex in a right-to-left, anterior-to-posterior and dorsal-to-ventral coordinate system. A gradient was defined as the sum of the difference between polar angle values of neighboring vertices multiplied by the differences between one vertex and neighboring vertices (vectors) divided by the sum of the squared Euclidean norm of the differences between vectors. If the number of connected vertices of a vertex is *n*, the gradient at vertex *k* is calculated as follows:

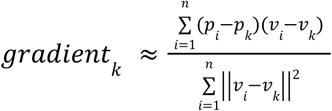

where *p* is the polar angle value of the vertex and *v* is a position vector of the vertex in a three-dimensional real coordinate space ℜ^3^ . V2 and V3 were subdivided into dorsal and ventral regions in accordance with the template created by Wang and colleagues ^53^ to allow for a more detailed group comparison.

### Statistical analyses

#### Adjustment for data quality and site effects

A parametric empirical Bayesian framework implemented in the *neuroCombat* Python library was employed before group comparison to rectify connectivity results for differences in data quality and site effects ^54^ . This framework incorporates age and sex as variables to be retained, while adjusting estimates for effects of no interest, including site, the averaged tSNR of V1 and the averaged tSNR of V2-3. The averaged tSNR was included since it is more representative than the tSNR of single vertices given that the connective field model is based on the weighted sum of multiple time courses from multiple vertices in V1.

#### Group comparison of connective field distributions

To ascertain whether the connective field estimates within each region of interest exhibit differential distributions across age groups, we employed bootstrapped Kruskal-Wallis tests written in custom Python code. The Kruskal-Wallis test allowed us to determine whether at least one group stochastically dominated at least one other group. Effect size was calculated as the square root of dividing the Kruskal-Wallis test statistic by the number of observations. The χ^2^ value was calculated by randomly resampling the subjects 10,000 times in all groups, using the same number of subjects in the original group and allowing duplication using SciPy’s *kruskal* function. Effect sizes of Kruskal-Wallis tests were calculated based on the following formula:

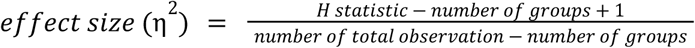

In the case of significant evidence for stochastic dominance, Mann-Whitney-U tests were performed vertex-wise for each pair of age groups. Mann-Whitney-U tests were carried out to compare pairs of groups using Scipy’s *mannwhitneyu* function. For paired group comparison, a bootstrap mean difference test was conducted by randomly resampling 10,000 times.

#### Modeling eccentricity and polar angle progression as a function of cortical distance

Linear regression models were fit to the individual progression of eccentricity across distance from the medial edge along V2 and V3. Then Wilcoxon signed-rank tests were run to confirm that the individual progression slopes differed significantly from zero, the equivalent of no change of eccentricity over distance. Absolute value functions centered on the border of V2 and V3 were fit to the progression of polar angle values and tested for statistical significance to model reversal of the progression at the border. To this end, the parameters of the absolute value function (parameters = a,b where y=a|x|+b) obtained from individual data were bootstrapped (n=10,000). The averaged polar angle values were then fitted to the reconstructed absolute value function using the bootstrapped parameters.

#### Comparison of connective field modeling results for measured and simulated data

We compared the results of the connective field model applied to the measured fMRI data in the V2 region (1) with the retinotopic values (eccentricity and polar angle) of Benson et al.’s template and (2) with the results of the connective field model applied to simulated fMRI data. For model comparison, we calculated the mean squared difference (MSD).

The following formula was employed to calculate the MSD values:

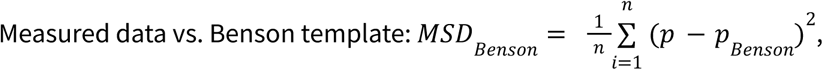

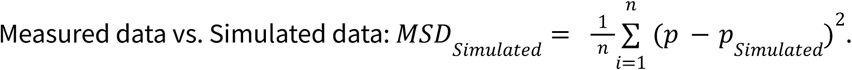

where 𝑛 is the number of vertices and 𝑝 is the connective field parameter value of interest for each vertex. Statistical differences between MSD values were calculated based on Wilcoxon signed-rank tests. Wilcoxon signed-rank tests were performed using Scipy’s *wilcoxon* function.

## Data availability

The data for this study are available after institutional registration through public links in the National Institute of Mental Health Data Archive (NDA). Prenatal and neonatal data are available at https://nda.nih.gov/edit_collection.html?id=3955. Adolescent and adult data are available at https://nda.nih.gov/edit_collection.html?id=2846.

## Code availability

All data analysis was performed with custom software using Python 3.11 and various third-party packages (matplotlib 3.8.3, numpy 1.26.3, pandas 2.2.0, scipy 1.12.0, and nibabel 5.2.0). Brain surfaces were visualized with nilearn (0.10.4). All code used for data analysis is available from GitHub (https://github.com/SkeideLab/ SPOT).

## Acknowledgements

This work was supported by the German Research Foundation (DFG Heisenberg Program Grant 433758790 awarded to M.A.S.), the Alexander von Humboldt Foundation (M.A.S. was appointed to the Henriette Herz Scouting Programme) and the Jacobs Foundation (Research Fellowship awarded to M.A.S.). Data were provided by the developing Human Connectome Project, KCL-Imperial-Oxford Consortium funded by the European Research Council under the European Union Seventh Framework Programme (FP/2007-2013) / ERC Grant Agreement no. [319456]. We are grateful to the families who generously supported this study.

## Author contributions

M.A.S. conceived the study. S.Y. and A.K. analyzed the data. S.Y. visualized the results with feedback from M.A.S.. S.Y. and M.A.S. wrote the manuscript. M.A.S. supervised the study.

## Ethics declarations

The authors declare no competing interests.

## Notes

### Competing Interest Statement

The authors have declared no competing interest.

### Summary of Updates

- analyzed a larger sample (N = 871) - confirmed generalizability of model performance by implementing a cross-validation approach

