## Supplementary Material for "Population connective field modeling reveals proto-retinotopic visual cortex organization in the prenatal human brain"

### Supplementary Figures S1-S11

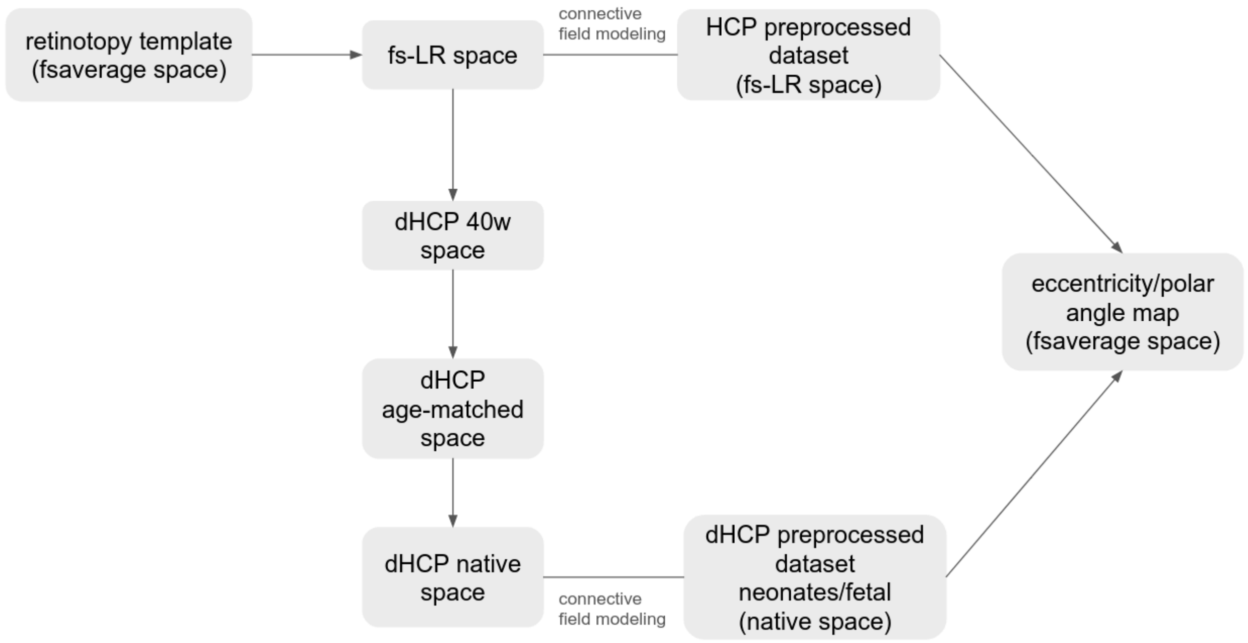


**Fig. S1 | Registration procedure.** A four-step registration procedure was applied to warp the retinotopy templates from Freesurfer's fsaverage space to native space, including (1) registration from fsaverage space to the space of the young adult template of the Human Connectome Project (HCP) in Freesurfer's fs-LR space, (2) registration from young adult space to the space of the 40-week template of the Developing Human Connectome Project (dHCP 40w) (see https://t.ly/AFwNT for both hemispheres), (3) registration from 40-week space to the space of an age-matched group template, and (4) registration from the age-matched group template space to native space.

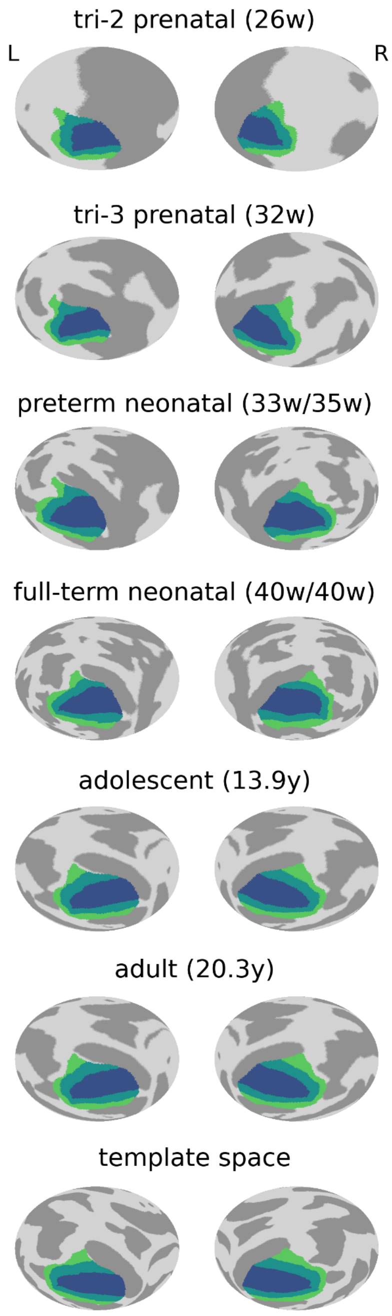


**Fig. S2 | Registration of individual structural images to the adult surface template containing the visual regions V1–V3.** Regions of interest (V1 = blue, V2 = teal, V3 = green) are shown on spherical surfaces in native space (L = left hemisphere, R = right hemisphere). Ages of participants are specified in brackets including birth age and scan age for preterm and full-term neonates (w = weeks, y = years).

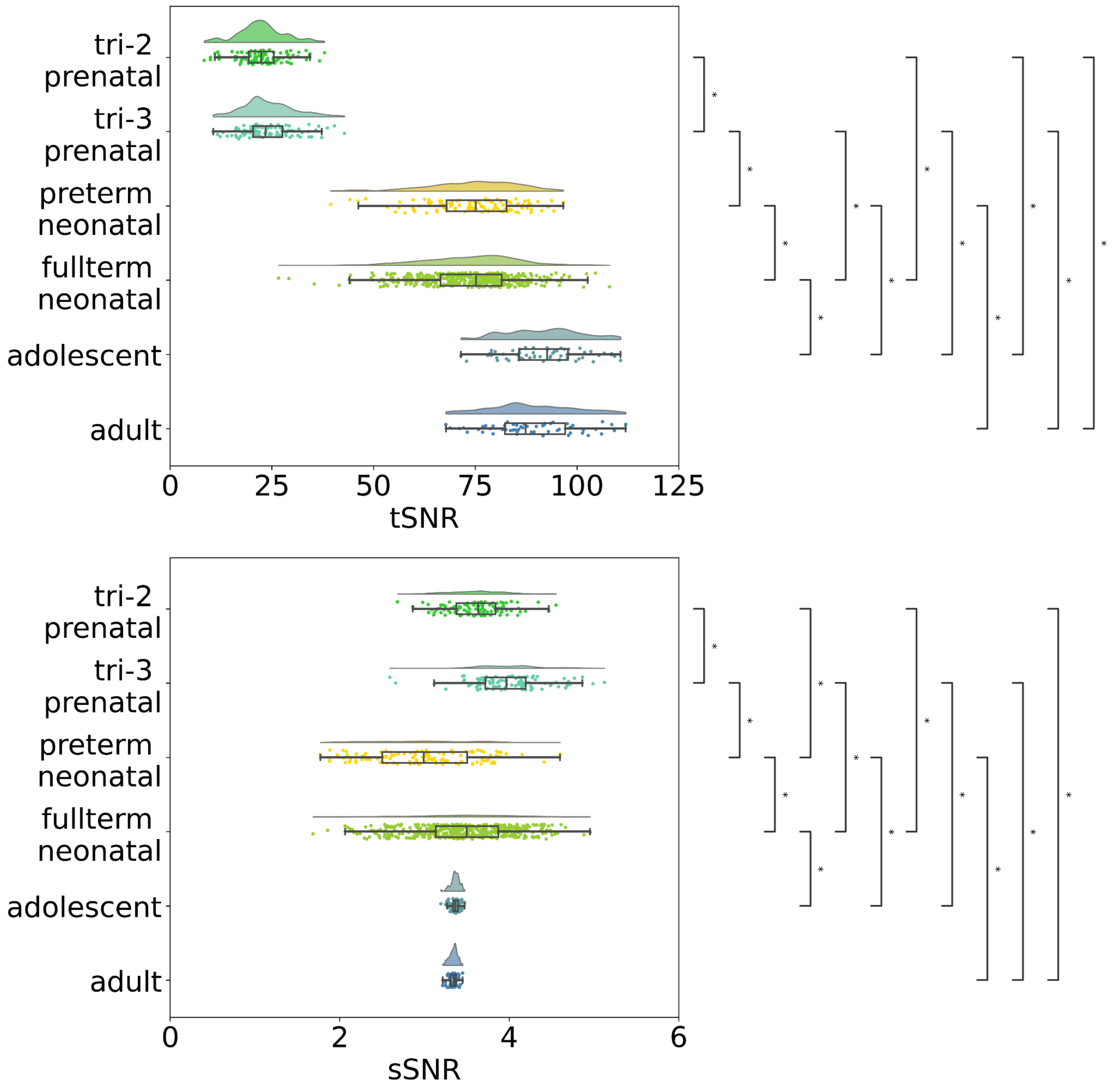


**Fig. S3 | Temporal and slice signal-to-noise ratio.** The x-axis depicts the temporal signal-to-noise ratio (tSNR, upper diagram) and the slice signal-to-noise ratio (sSNR, lower diagram) of the resting-state fMRI data. The y-axis depicts the different age groups. Curves illustrate probability density while lines illustrate the lengths of the distributions. Box plots visualize upper and lower quartiles. Asterisks indicate statistically significant differences at a bootstrapped threshold of $p < 0.001$ (Mann-Whitney-U tests).

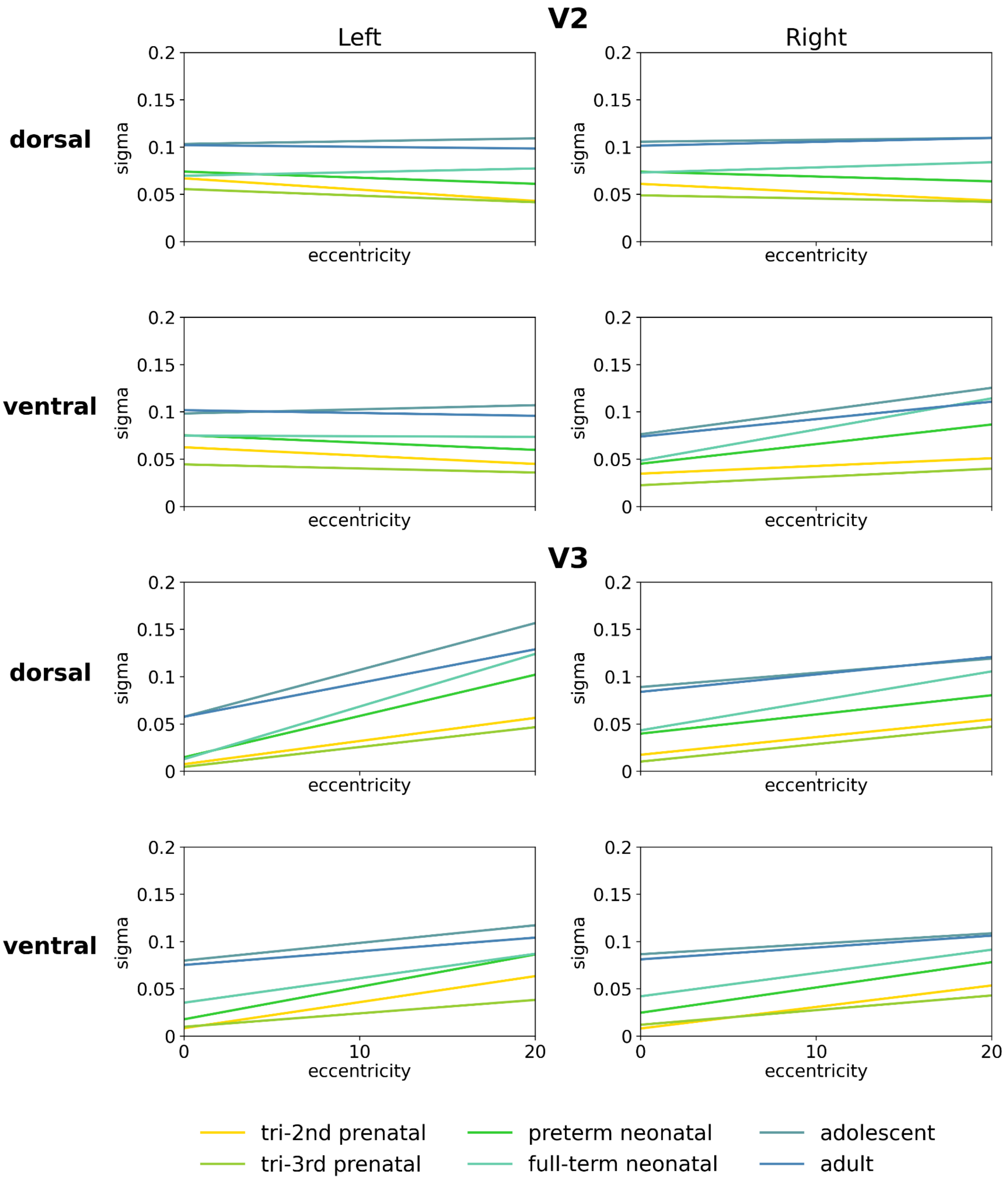


**Fig. S4 | Associations between eccentricity and connective field size.** The x-axis shows eccentricity (degrees visual angle) and the y-axis shows connective field size (sigma). Lines represent linear regression fits to group-average values. These values are not shown for better visibility. Lines are color-coded by group (see legend).

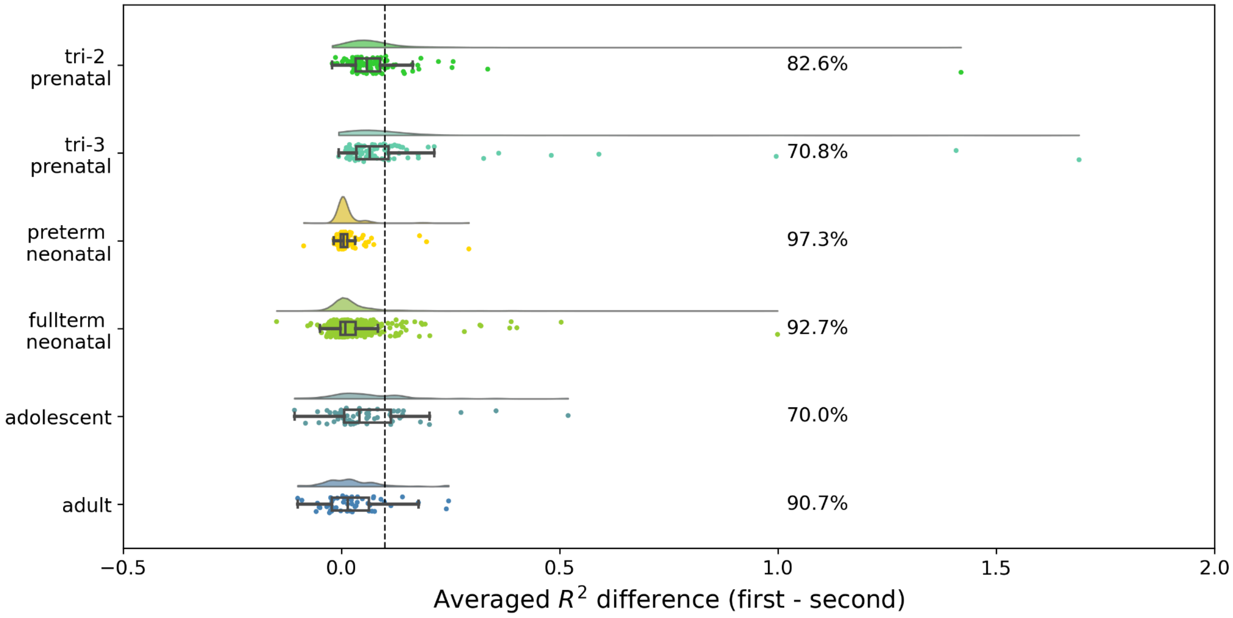


**Fig. S5 | Split-half cross-validation results.** The x-axis depicts each age group and the y-axis depicts the individual differences in variance explained ($\Delta R^2 = R_1^2 - R_2^2|$) between the first and the second half of the hemodynamic time series, averaged across regions of interest and hemispheres. The vertical dashed line indicates the chosen reliability threshold ($\Delta R^2 = 0.1$). Values to the left of this line correspond to subjects for which model generalization is considered as reliable. Percentages indicate the proportion of subjects in each group passing this threshold. Boxplots and density distributions summarize the spread of $\Delta R^2$ values within each group.

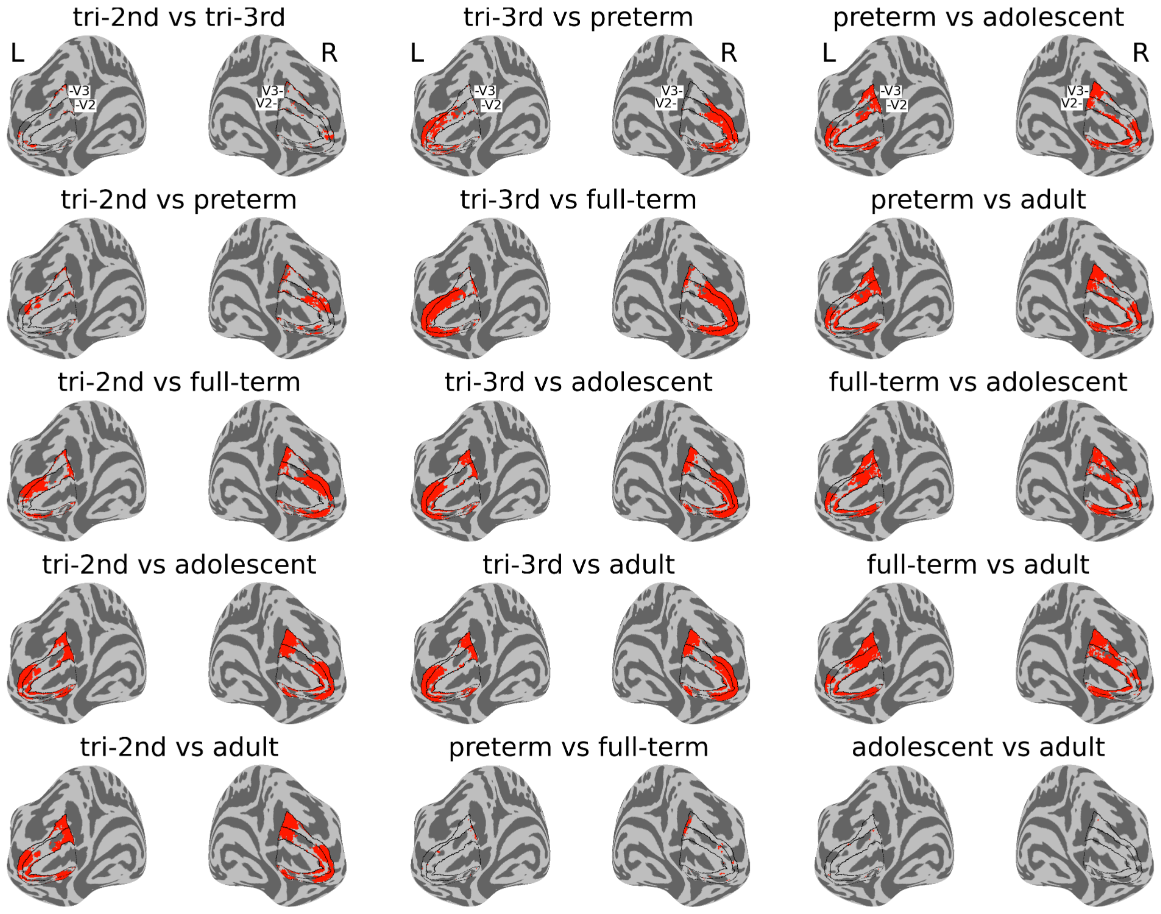


**Fig. S6 | Group differences in eccentricity.** Red vertices demonstrate pairwise group differences in eccentricity surface maps (family-wise error corrected at $p < 0.001$, mean difference bootstrapping at $p < 0.001$, Mann-Whitney-U tests). These maps were adjusted for effects of site and data quality. tri-3rd = third trimester prenatal. tri-2nd = second trimester prenatal. preterm = preterm neonatal. full-term = full-term neonatal. L = left hemisphere, R = right hemisphere. Black lines demarcate V1, V2, and V3.

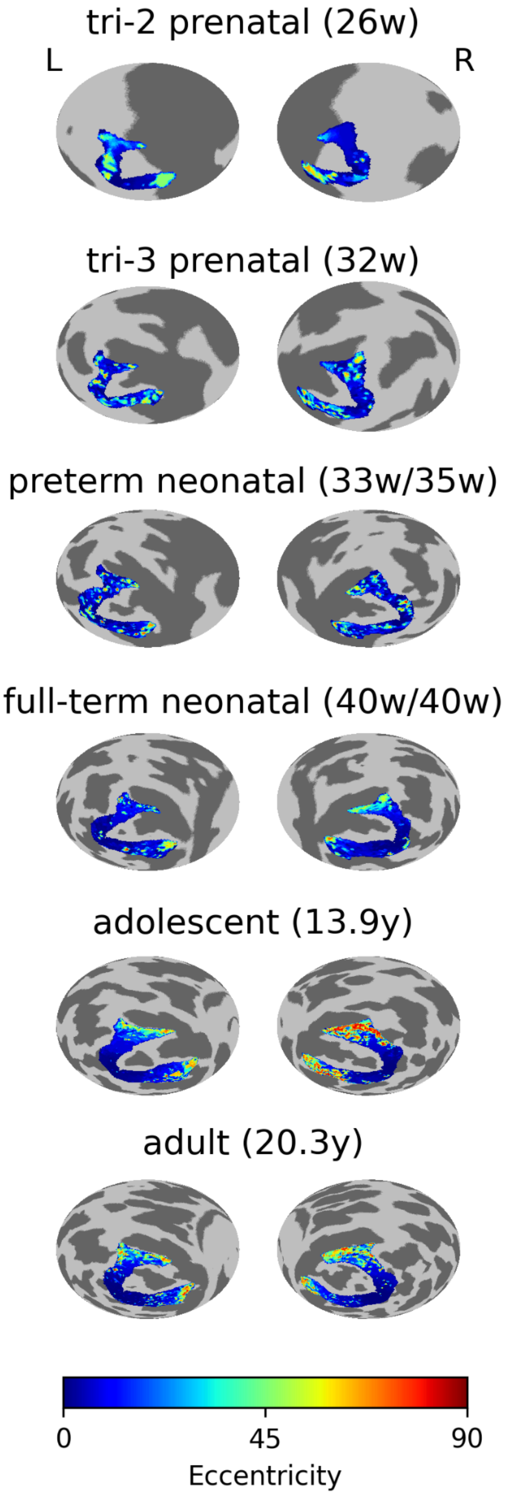


**Fig. S7 | Individual eccentricity maps.** L = left hemisphere, R = right hemisphere. Ages of participants are specified in brackets including birth age and scan age for preterm and full-term neonates (w = weeks, y = years). The color bar indicates eccentricity estimates (degrees from the fovea). Regions of interest include V2 and V3.

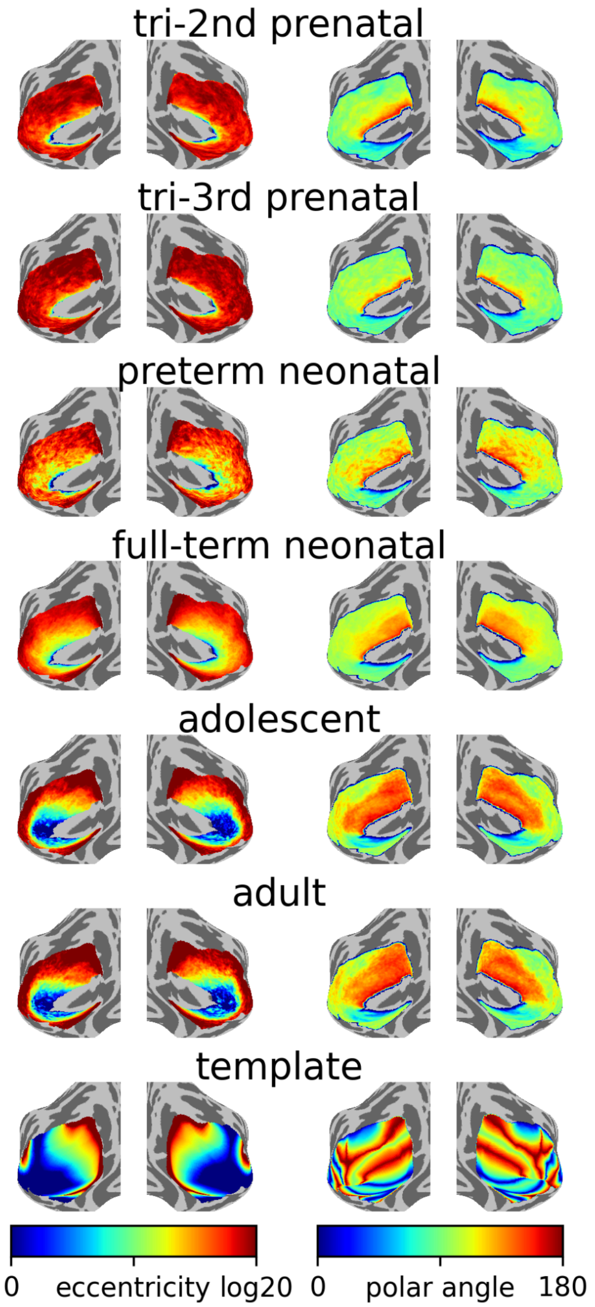


**Fig. S8 | Eccentricity and polar angle maps without predefined extrastriate regions of interest.** Group-averaged eccentricity values (two left columns) and polar angle values (two right columns). The left color bar indicates log-transformed eccentricity and the right color bar indicates polar angle (both in degrees of visual angle). L = left hemisphere, R = right hemisphere.

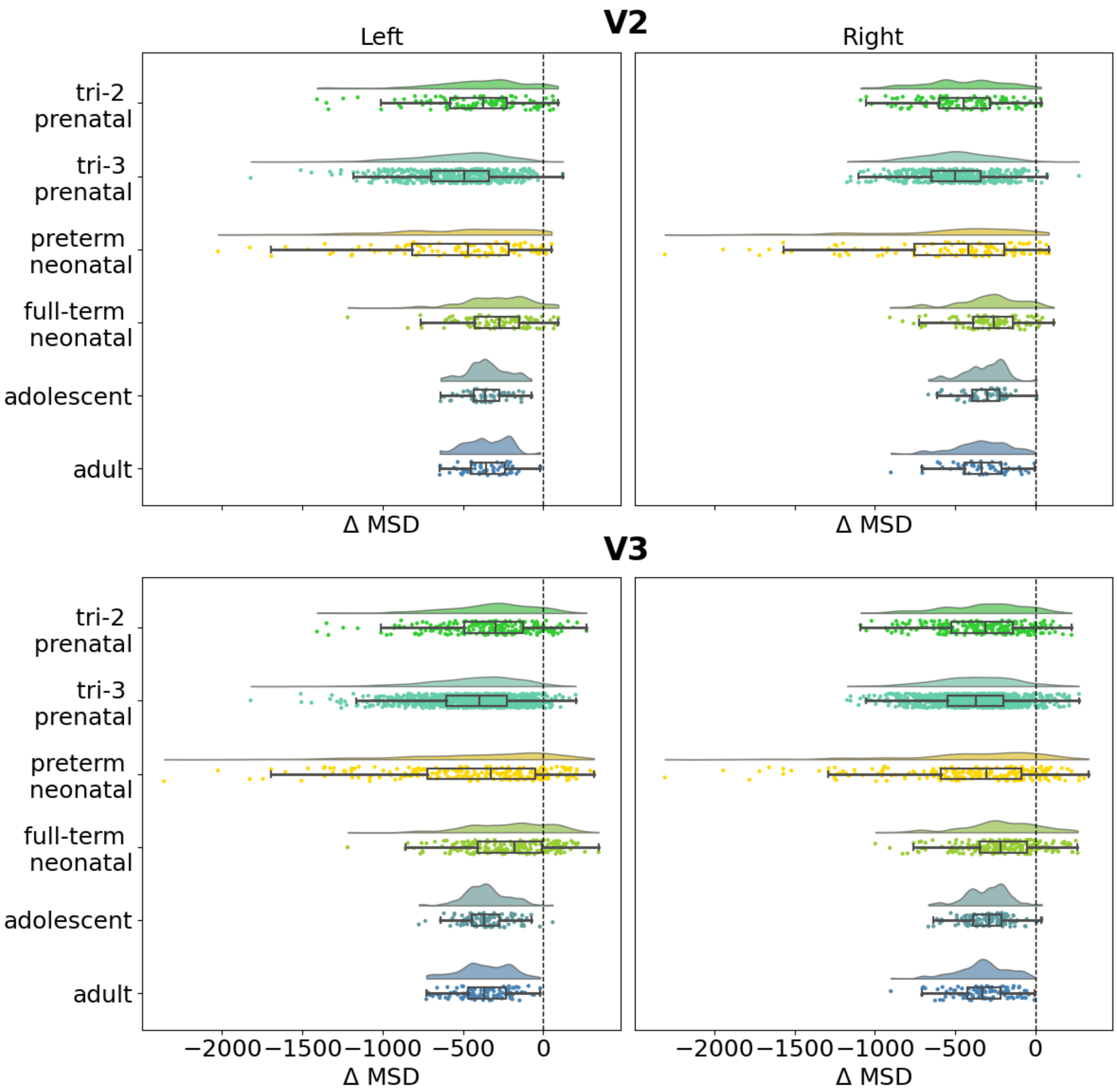


**Fig. S9 | Differences between empirical and simulated eccentricity estimates.** The x-axis depicts the mean-squared difference between the eccentricity template data and the empirical estimates subtracted by the mean-squared difference between the eccentricity template data and estimates obtained from random simulated hemodynamic signals (Δ MSD). Negative values on the x-axis visualize that eccentricity estimates obtained from the empirical fMRI data are more similar to the template than random simulated data. The y-axis depicts the different age groups. Curves illustrate probability density while solid lines illustrate the lengths of the distributions. Box plots visualize upper and lower quartiles. Dashed vertical lines extend the zero point of the x-axis.

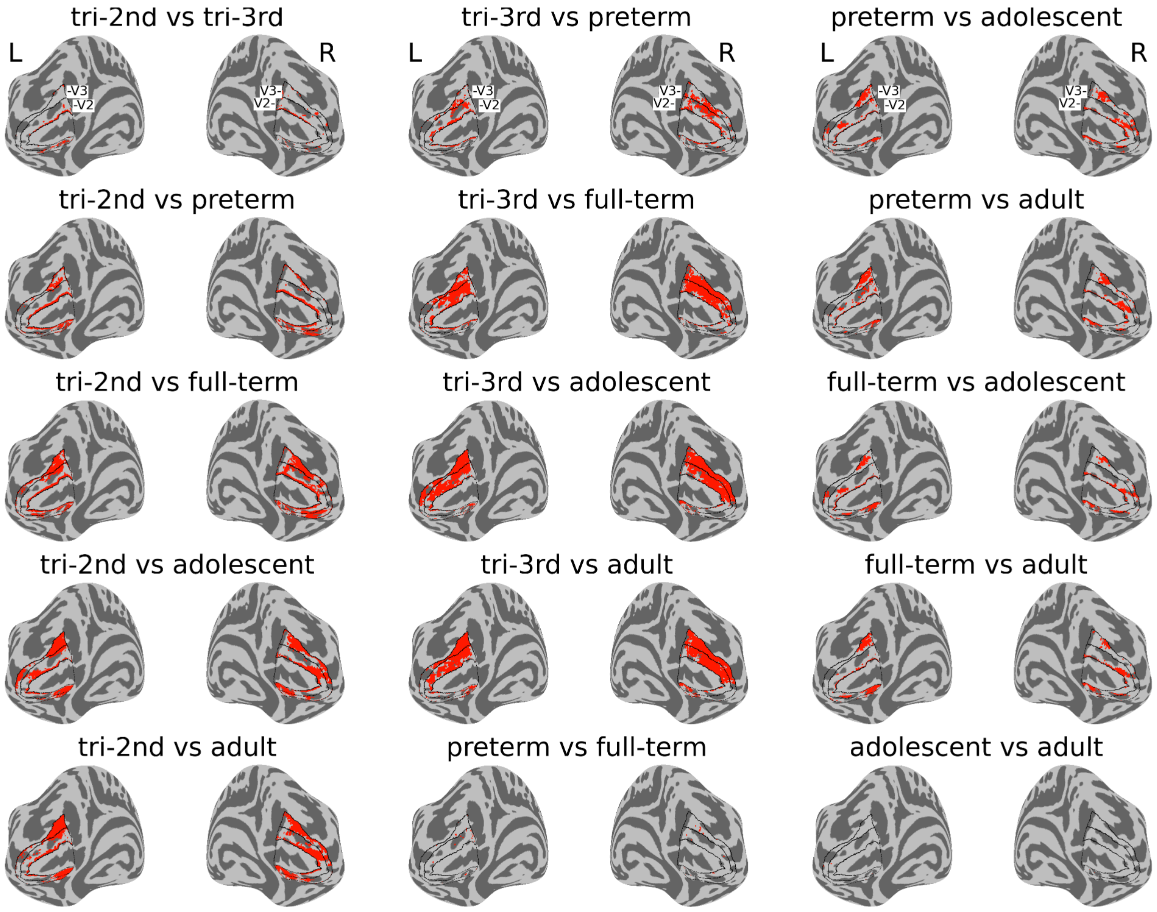


**Fig. S10 | Group differences in polar angle.** Red vertices demonstrate pair-wise group differences in polar angle surface maps (family-wise error corrected at p < 0.001, mean difference bootstrapping at p < 0.001, Mann-Whitney-U tests). These maps were adjusted for effects of site and data quality. tri-3rd = third trimester prenatal. tri-2nd = second trimester prenatal. preterm = preterm neonatal. full-term = full-term neonatal. L = left hemisphere, R = right hemisphere. Black lines demarcate V1, V2, and V3.

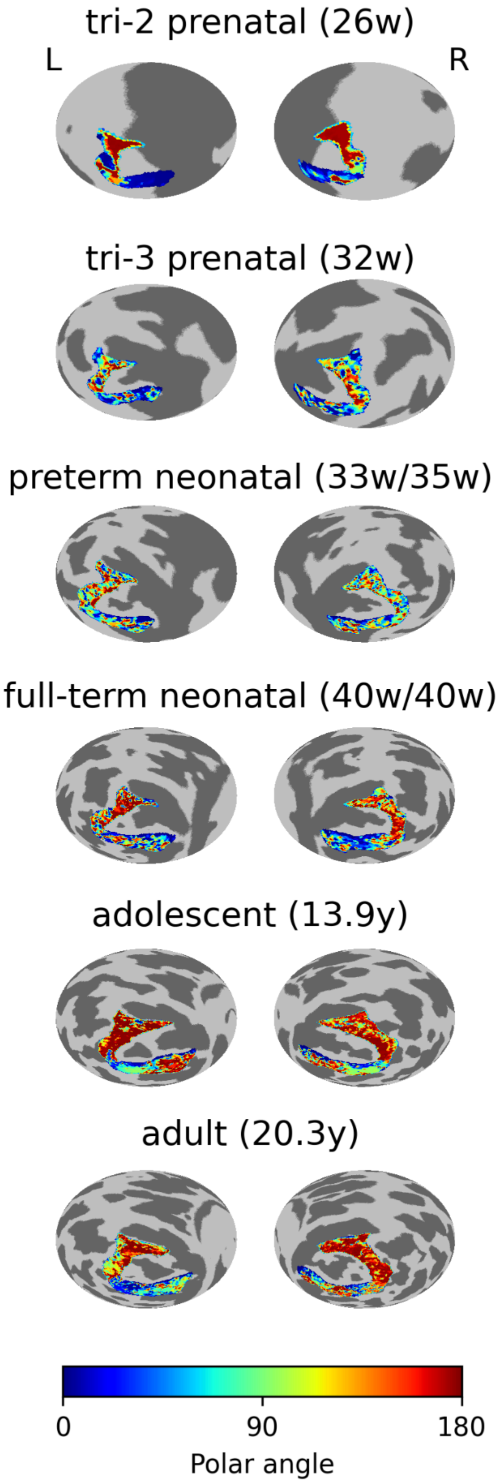


**Fig. S11 | Individual polar angle maps.** L = left hemisphere, R = right hemisphere. Ages of participants are specified in brackets including birth age and scan age for preterm and full-term neonates (w = weeks, y = years). The color bar indicates polar angle estimates (angular distance from the vertical median in degrees). Regions of interest include V2 and V3.

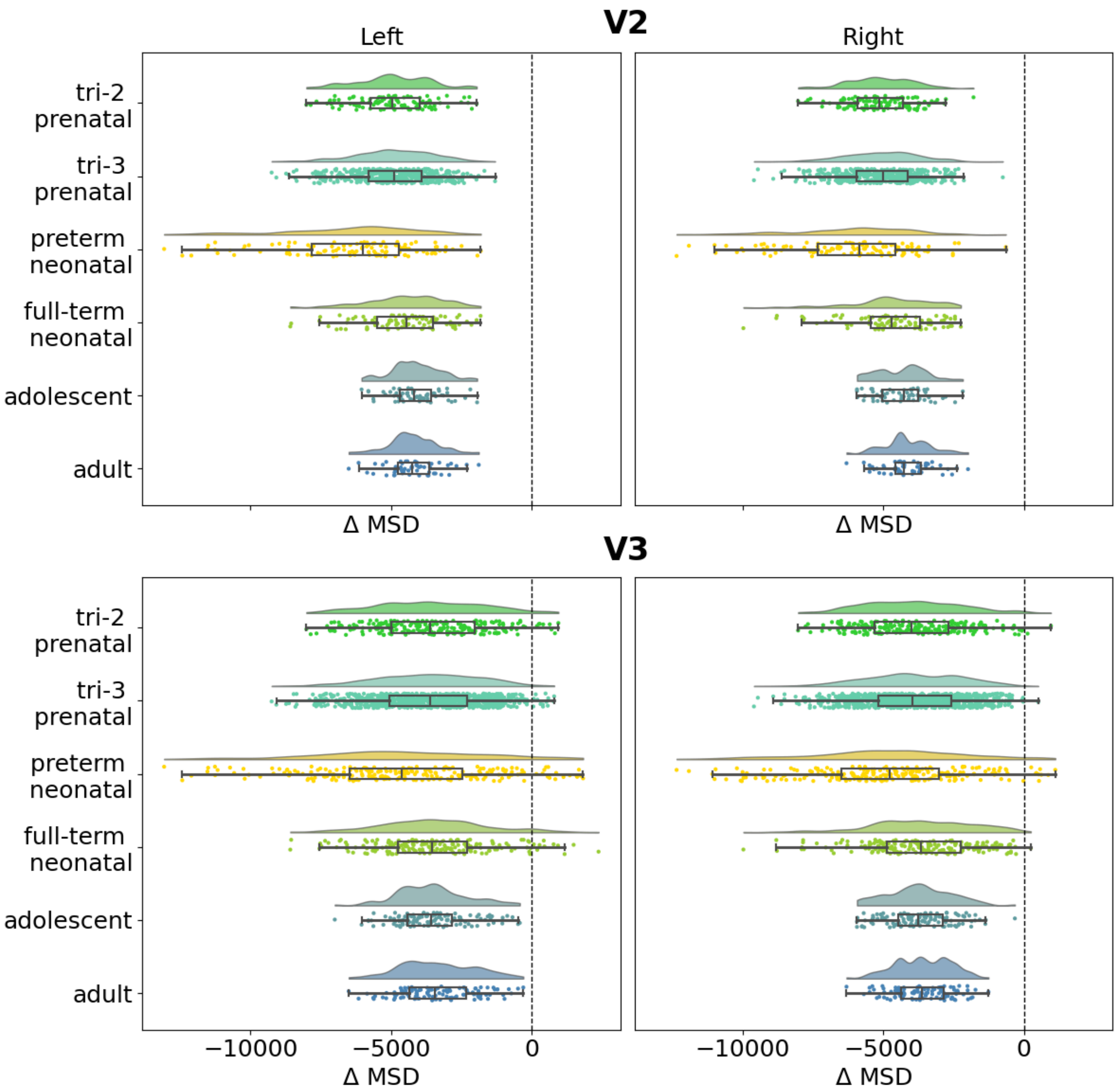


**Fig. S12 | Differences between empirical and simulated polar angle estimates.** The x-axis depicts the mean-squared difference between the polar angle template data and the empirical estimates subtracted by the mean-squared difference between the polar angle template data and estimates obtained from random simulated hemodynamic signals (Δ MSD). Negative values on the x-axis visualize that polar angle estimates obtained from the empirical fMRI data are more similar to the template than random simulated data. The y-axis depicts the different age groups. Curves illustrate probability density while solid lines illustrate the lengths of the distributions. Box plots visualize upper and lower quartiles. Dashed vertical lines extend the zero point of the x-axis.
